# Semi-field evaluation of the space spray efficacy of Fludora Co-Max EW against wild insecticide-resistant *Aedes aegypti* and *Culex quinquefasciatus* mosquito populations from Abidjan, Côte d’Ivoire

**DOI:** 10.1101/2022.05.03.490391

**Authors:** Julien Z. B. Zahouli, Jean-Denis Dibo, Fofana Diakaridia, Laurence Yao, Sarah D. Souza, Sebastian Horstmann, Benjamin G. Koudou

**Affiliations:** Centre Suisse de Recherches Scientifiques en Côte d’Ivoire, Abidjan, Côte d’Ivoire; Centre d’Entomologie Médicale et Vétérinaire, Université Alassane Ouattara, Bouaké, Côte d’Ivoire; Unité de Formation et de Recherche Sciences de la Nature, Université Nangui-Abrogoua, Abidjan, Côte d’Ivoire; Institut National d’Hygiène Publique, Ministère de la Santé et de l’Hygiène Publique, Abidjan, Côte d’Ivoire; Environmental Science, Bayer AG Crop Science Division, Bayer, Nairobi, Kenya; Environmental Science, Bayer AG Crop Science Division, Bayer, Monheim, Germany

## Abstract

**Background:** Space spraying of insecticides is still an important mean of controlling *Aedes* and *Culex* mosquitoes and arboviral diseases. This study evaluated the space spray efficacy of Fludora Co-Max EW (a combination of flupyradifurone and transfluthrin, with Film Forming Aqueous Spray Technology (FFAST)) against wild, insecticide-resistant *Aedes aegypti* and *Culex quinquefasciatus* populations from Abidjan, Côte d’Ivoire, against K-Othrine EC (deltamethrin-only product), through small-scale field trials.

**Methodology:** Wild *Ae. aegypti* and *Cx. quinquefasciatus* mosquito larvae were collected in Abidjan, Côte d’Ivoire from August to December 2020. Mosquito larvae were reared until adult stage. Emerged adult females were tested against Fludora Co-Max EW and K-Othrine EC using ultra-low volume cold fogging (ULV) and thermal fogging (TF) both outdoors and indoors in Agboville, Côte d’Ivoire. Cages containing 20 mosquitoes each were placed at 10, 25, 50, 75 and 100 m from the spraying line for outdoor spraying, and at ceiling, mid-height and floor levels for indoor house spraying. Knockdown and mortality were recorded at each checkpoint and compared by treatments.

**Principal findings:** Overall, Fludora Co-Max EW induced significantly higher knockdown and mortality effects in the wild insecticide-resistant *Ae. aegypti* and *Cx. quinquefasciatus* compared with K-Othrine EC. With both species, Fludora Co-Max EW mortality rates were above 80% (up to 100%) for outdoor ULV spray at each distance checkpoint (i.e. 10 to 100 m), and 100% for indoor ULV and TF sprays at all level checkpoints (i.e. ceiling, mid-height and floor). K-Othrine EC induced high mortality indoors (97.9-100%), whereas outdoor mortality rapidly declined in *Ae. aegypti* from 96.7% to 36.7% with ULV, and 85.0% to 38.3% with TF, from 10 to 100 m. For outdoor Fludora Co-Max EW spray, ULV showed both higher knockdown and killing performance *Ae. aegypti* and *Cx. quinquefasciatus* compared with TF. Fludora Co-Max EW performed better against *Cx. quinquefasciatus* compared with *Ae. aegypti*.

**Conclusion/significance:** Fludora Co-Max EW induced high mortality and knockdown effects against wild insecticide-resistant *Ae. aegypti* and *Cx. quinquefasciatus* Abidjan strains and performed better than K-Othrine EC. The presence of flupyradifurone and transfluthrin (with new and independent modes of action) and FFAST technology in the current Fludora Co-Max EW formulation appears to have broadened its killing capacity. Fludora Co-Max EW is thus an effective adulticide and may be a useful tool for *Aedes* and *Culex* mosquito and arbovirus control in endemic areas.

**Author Summary:** Space spraying of insecticides is an important tool to control *Aedes* and *Culex* mosquitoes and prevent the viral diseases (i.e. dengue, yellow fever, etc.) that they transmit. We studied the efficacy of the product Fludora Co-Max EW (a new space spray insecticide) against adult wild insecticide-resistant populations of *Aedes aegypti* and *Culex quinquefasciatus* mosquitoes from Abidjan, Côte d’Ivoire. We compared Fludora Co-Max EW knockdown and mortality effects in these mosquitoes with the local insecticide K-Othrine EC using ultra-low volume (ULV) and thermal fogging (TF) spraying outdoors and indoors. The product Fludora Co-Max EW induced high rates of knockdown and mortality (i.e. 80-100%) in these wild insecticide-resistant mosquitoes and performed better than the product K-Othrine EC. Additionally, ULV sprays of Fludora Co-Max EW demonstrated higher knockdown and killing efficacy at larger distances (i.e. up to 100 m) compared with TF. The higher efficacy of Fludora Co-Max EW may be due to the interaction of two unrelated insecticides, flupyradifurone and transfluthrin, in combination with Film Forming Aqueous Spray Technology (FFAST). Fludora Co-Max EW therefore appears to be an effective and useful tool to control adult populations of wild insecticide-resistant *Aedes* and *Culex* mosquitoes and may be recommended for preventing related mosquito-transmitted viral diseases.

## Introduction

*Aedes aegypti* and *Culex quinquefasciatus* mosquitoes are the primary vectors of arboviruses (e.g. dengue, yellow fever, chikungunya, Zika, Rift valley fever and West Nile virus) transmitted to humans and livestock worldwide [1]. For instance, over 3.9 billion people are at risk of dengue fever, with about 390 million infections per year, resulting in over 100 million symptomatic cases and 10,000 deaths [2]. Arboviruses and their *Aedes* vectors are spreading rapidly worldwide from their African origin [3], due to accelerated urbanisation, international trade, mobility and climate change [4, 5]. In Africa, arboviruses are threatening over 831 million people, which is 70% of the continental population [6–8], and Rift Valley fever infections have caused increased abortions, stillbirths and mortality among domestic animals with considerable economic losses [9]. Because there are no vaccines and no specific drugs against most of these arboviruses, prevention is mainly based on an effective surveillance and vector control [1]. Space spraying of insecticides is still an important mean of controlling *Aedes* and *Culex* mosquitoes and preventing arboviral outbreaks [10].

As in many African countries, dengue has become a major public health concern in Côte d’Ivoire, with emergence of multiple outbreaks often coupled with yellow fever co-infections [11, 12]. The majority (over 80%) of these outbreaks have occurred in the city of Abidjan (∼6 million inhabitants) [11, 12]. The outbreak responses, conducted by the national arbovirus control programme of the Ministry of Health (MoH) and based on space sprays of insecticides (e.g. deltamethrin and chlorpyriphos-ethyl), have often resulted in limited and short-term impacts on the *Aedes* vector control: *Ae. aegypti* populations recover quickly and arboviruses re-emerge massively once campaigns are over [11, 12]. The livestock was infected by Rift Valley fever virus in Côte d’Ivoire [3]. Indeed, the local *Ae. aegypti* and *Cx. quinquefasciatus* populations are highly abundant [14, 15], and resistant to many insecticides (e.g. deltamethrin, permethrin, lamdacyhalothrin, propoxur and fenitrothion) used for their control [16–18].

Given the present spread of arboviruses and insecticide resistance in *Aedes* arbovirus vectors worldwide [19–21] and from West Africa [22–24], there is a pressing need to develop effective vector tools and strategies to prevent arboviral epidemics. Sustainable and rational deployment of an effective and eco-friendly tool for outdoor and indoor vector control within insecticide resistance management programmes can reduce local vector densities, save lives and financial resources, and improve human, animal and environmental health. It is therefore important to develop new vector control tools incorporating a novel insecticide molecule or a combination of insecticide molecules with different modes of action, as part of insecticide resistance management. The candidate product to be evaluated in the current study is Fludora Co-Max EW, a new mixture of formulations developed by Bayer for outdoor and indoor space spray. The Fludora Co-Max EW formulation is a combination of two insecticides, flupyradifurone and transfluthrin, and is primarily based on water emulsion with built-in anti-evaporant technology, Film Forming Aqueous Spray Technology (FFAST), for longer space impaction efficiency.

This small-scale field study aimed to evaluate the space spray efficacy of Fludora Co-Max EW against adult females of natural and insecticide-resistant populations of *Ae. aegypti* and *Cx. quinquefasciatus* in comparison with K-Othrine EC (only deltamethrin product) in Abidjan, Côte d’Ivoire. As Fludora Co-Max EW s built on a combination of two new insecticides with unilateral modes of action and FFAST technology, it is expected to kill the wild pyrethroid-resistant *Ae. aegypti* and *Cx. quinquefasciatus* Abidjan strains. The results will show the potential of Fludora Co-Max EW for contributing to the management of insecticide resistance in *Aedes* and *Culex* mosquitoes and providing sustainable, cost-effective and environmentally friendly solutions for the control of arboviral diseases.

## Methodology

### Mosquitoes

*Ae. aegypti* and *Cx. quinquefasciatus* mosquitoes from Abidjan, Côte d’Ivoire were used in this trial. Both mosquito Abidjan strains have shown varied resistance to insecticides; ranging from probable resistance and confirmed resistant to the insecticides (e.g. K-Othrine) usually used for their control in Côte d’Ivoire [16–18]. Recent studies detected three knockdown resistance (*kdr*) mutations (V410L, V1016I and F1534C) in the wild *Ae. aegypti* Abidjan populations, with low frequencies for Leu 410 (0.28) and Ile1016 (0.32) alleles and high frequency for Cys1534 allele (0.96) [18]. *Aedes* and *Culex* mosquito immatures (i.e. larvae and pupae) were collected in their common breeding sites (e.g. tyres, tins cans, discarded containers, puddles, gutters, etc.) in Abidjan (5° 20’ 11” North, 4° 01’ 36” West). Collected mosquito immatures were transported by air-conditioned car to our insectary in Agboville (100 km from Abidjan), Côte d’Ivoire for rearing until adult stage and testing. They were reared until adult stage under standard conditions (27 ± 2 °C, 80 ± 10% relative humidity (RH), 12: 12 hour light: dark). Emerged *Ae. aegypti* and *Cx. quinquefasciatus* adults were fed with a 10% sugar solution soaked cotton wool and maintained under similar laboratory conditions. Unfed females aged of 3-5 days of each species were caged by 20 specimens per evaluation cage used in the trials.

### Insecticide formulations and dilution

The candidate Fludora Co-Max EW 78.8 and the local standard product K-Othrine EC 25 water-based space spray concentrates are specifically designed for dilution in water only. Fludora Co-Max EW is a combination of two insecticides, flupyradifurone (Flupyradifurone: 26.3 g/L) and transfluthrin (Transfluthrin: 52.5 g/L) with unrelated modes of action and features FFAST technology for longer space impaction efficiency, while K-Othrine EC is deltamethrin (Deltamethrin: 25 g/L) only product.

In this trial, there were three study arms (candidate product: Fludora Co-Max EW 78.8, positive reference control: K-Othrine EC 25, and negative control: water without insecticide) per application method (ultra-low volume cold fogging (ULV) and thermal fogging (TF) outdoors and indoors). Therefore, the insecticide products were diluted with the suitable volume of water according to the equipment manufacturer’s recommendations for the output rates required (S1 Fig). Three replicates were performed per concentration and per insecticide formulation, including the negative control, and fresh dilution quantities required for immediate usage were prepared for each new replicate. The dilution rates of each formulation (i.e. Fludora Co-Max EW 78.8 and K-Othrine EC 25) to be used for the experiments were:1:10, 1:50, 1:100 and 1:100 for outdoor ULV, indoor ULV, outdoor TF and indoor TF, respectively.

### Spray equipment

As standard practice, all application equipment was calibrated to determine output rates and actual volumes required in order to reach the product target application rates. A vehicle-mounted sprayer was used for outdoor spraying and a hand-held sprayer for indoor sprays. As no coated slides or droplet catching machines were available, the manufacturer guidelines for machine calibration were followed to take in account the droplet size. A nozzle for the correct volume medium diameter (VMD) size for the space spray trial was expected to be between 25 and 30 µm. The discharge rate was pre-calibrated in a separate spraying exercise. An experienced operator conducted the fogging, moving along the spray line at a standard pace.

### Trial area and procedures

This Phase II semi-field study was carried out through four space spray trial types: outdoor ULV, outdoor TF, indoor ULV and indoor TF (S2 Fig). All trials were carried out in Agboville (5° 55’ 41” North, 4° 13’ 01” West). Outdoor trials were conducted in an open area with no vegetation, namely “Place Bédié” (250 m x 150 m). Indoor trials were performed in an empty room of a house, with an approximate volume of 126 m^3^ (8.5 m long x 5.3 width x 2.8 height). The trials were designed and carried out according to the current standard World Health Organization (WHO) guidelines for outdoor and indoor space spray applications of insecticides for the control of vectors and public health pests [25, 26]:

#### Outdoor fogging

At the “Place Bédié”, a 200 m straight transect was designated on one side of this rectangular site as a spray line to be followed by the spraying vehicle. Perpendicularly to and downwind from the spray line, 1.5 m tall poles were set at five distance checkpoints of 10, 25, 50, 75 and 100 m (S3 Fig). A checkpoint evaluation cage with 20 mosquitoes was placed at the top of each pole. The cages were cylindrical and constructed of fine mesh fabric (nylon) with wire frame support (diameter 10 cm x height 15 cm x tapping cover 10 cm) (S4 Fig). The mesh sizes were around 1.2 mm, to allow a throughput of spray-droplets. The vehicle-mounted spray machine was used for the tests. The wind direction should be perpendicular to the vehicle travel line, and was assessed using a long low-weight tissue (e.g. toilet paper). After spraying, tested mosquitoes were left in the test area for 15-minute exposure. Spraying was conducted in the morning before the sunrise and evening after sunset. The ambient temperature and RH were monitored and recorded before and after each trial. The meteorological conditions (25 ± 1 °C; 80 ± 10% RH) were quite optimal for the space spraying.

#### Indoor fogging

The room selected for indoor spraying was ventilated and adequately decontaminated. A total of 10 checkpoint evaluation cages per mosquito species (20 mosquitoes/cage) were placed 25 cm from each corner at ceiling and floor levels and two at mid-height (approximatively 1.5 m near the centre of the room (S5 Fig). Windows were closed prior to the application. An experienced technician delivered the spray from the front door (i.e. inside-out treatment), or through an open window with the hand-held sprayer nozzle directed towards the centre of the room as adequate dispersal of the insecticide droplets were achieved. After the application, external doors and windows were closed during the mosquitoes’ 20-minute exposure period. The environmental conditions in the test room were recorded before and after each trial. The weather (25 ± 1°C, 80 ± 10% RH) was fair during the test.

#### Evaluation

After the specified exposure period, knockdown was immediately recorded at the time of collection. Then, tested mosquitoes were rapidly transferred into clean holding containers (plastic cups) covered with mesh cloth using an appropriate aspirator and brought back to the field station. Mosquitoes knocked out were recorded every 3 and 2 minutes alternatively during 60 minutes, thus allowing to record knockdown at 0, 10, 30, 40, 50 and 60 minutes. The tested mosquitoes were provided with a 10% sucrose cotton wool pad, and transferred to the laboratory. In the laboratory, they were held at standard conditions (27 ± 2 °C, 80 ± 10% RH, 12:12 hour light: dark) for 24 hours. The 24-hour mortality was recorded.

#### Test validity

The mortality rates in the concurrent negative control cages were less than 5%, thus the results from the treated samples were directly accepted.

### Side effect assessment

We assessed the possible side effects (e.g. itching, dizziness or nose running) among the sprayers and the occupants of the sprayed house through the interviews.

### Statistical analysis

Raw data were double entered into Excel using double key entry, and checked for outliers. The knockdown rate was expressed as the percentage of specimens knocked out (i.e. numerator) among mosquitoes tested (i.e. denominator). The mortality rate was calculated as the percentage of dead specimens (i.e. numerator) among mosquitoes exposed (i.e. denominator) to treatment for each insecticide dose. The mortality rates (mean ± standard error) were compared using a one-way analysis of variance (ANOVA), followed by Bonferroni’s correction. The data were analysed by Kruskal-Wallis test only when they showed significant deviations from normality (i.e. when Shapiro-Wilk W test for normal data was significant). A significance level of 5% was set for statistical testing. All statistical analyses were conducted using Stata version 16.0 (Stata Corporation; College Station, TX, United States of America).

### Ethics statement

The study protocol was approved by the National Ethics Committee for Research of Ministry of Health and public Hygiene in Côte d’Ivoire (N/Ref: 113-20/MSHP/CNESVS-km) and the health district and administrative authorities of Abidjan and Agboville. Signed informed consent was obtained from owners and/or residents before conducting larval collections and spraying in or near a residence.

## Results

### Outdoor ULV efficacy

#### Knockdown

At 0 minutes post-exposure, the knockdown rates of Fludora Co-Max EW and K-Othrine EC outdoor ULV space sprays from 10 to 100 m were of 100.0-85.0% and 83.3-28.3% in *Ae. aegypti*; and 100.0-3.3%, and 96.7-38.3% in *Cx. quinquefasciatus*, respectively (S1 Table). At 60 minutes after exposure, Fludora Co-Max EW and K-Othrine EC induced respective knockdown rates of 100.0-95.0% and 91.7-30.0% in *Ae. aegypti*, and 100.0-95.0% and 100.0-78.3% in *Cx. quinquefasciatus* 10 to 100 m (S1 Table). Thus, Fludora Co-Max EW effected faster and higher knockdown up to a longer distance compared to K-Othrine EC in both species.

#### Mortality

Outdoor ULV space sprays of Fludora Co-Max EW and K-Othrine EC resulted in respective overall mortality rates of 92.3 ± 2.1% and 66.7 ± 5.5% in *Ae. aegypti*, and of 99.7 ± 0.3% and 95.3 ± 1.6% in *Cx. quinquefasciatus* (S2 Table). Fludora Co-Max EW mortality rates were significantly higher compared with K-Othrine EC in *Ae. aegypti* (χ^2^ = 11.13; df = 1; p < 0.05) and *Cx. quinquefasciatus* (χ^2^ = 6.30; df = 1; p < 0.05). Additionally, Fludora Co-Max EW mortality rates were very high and above 90% in both mosquito species, whereas K-Othrine EC mortality was higher than 90% only in *Cx. quinquefasciatus*.

The mortality rates varied substantially depending on the distance checkpoint (Figure 1).

**Figure 1:**
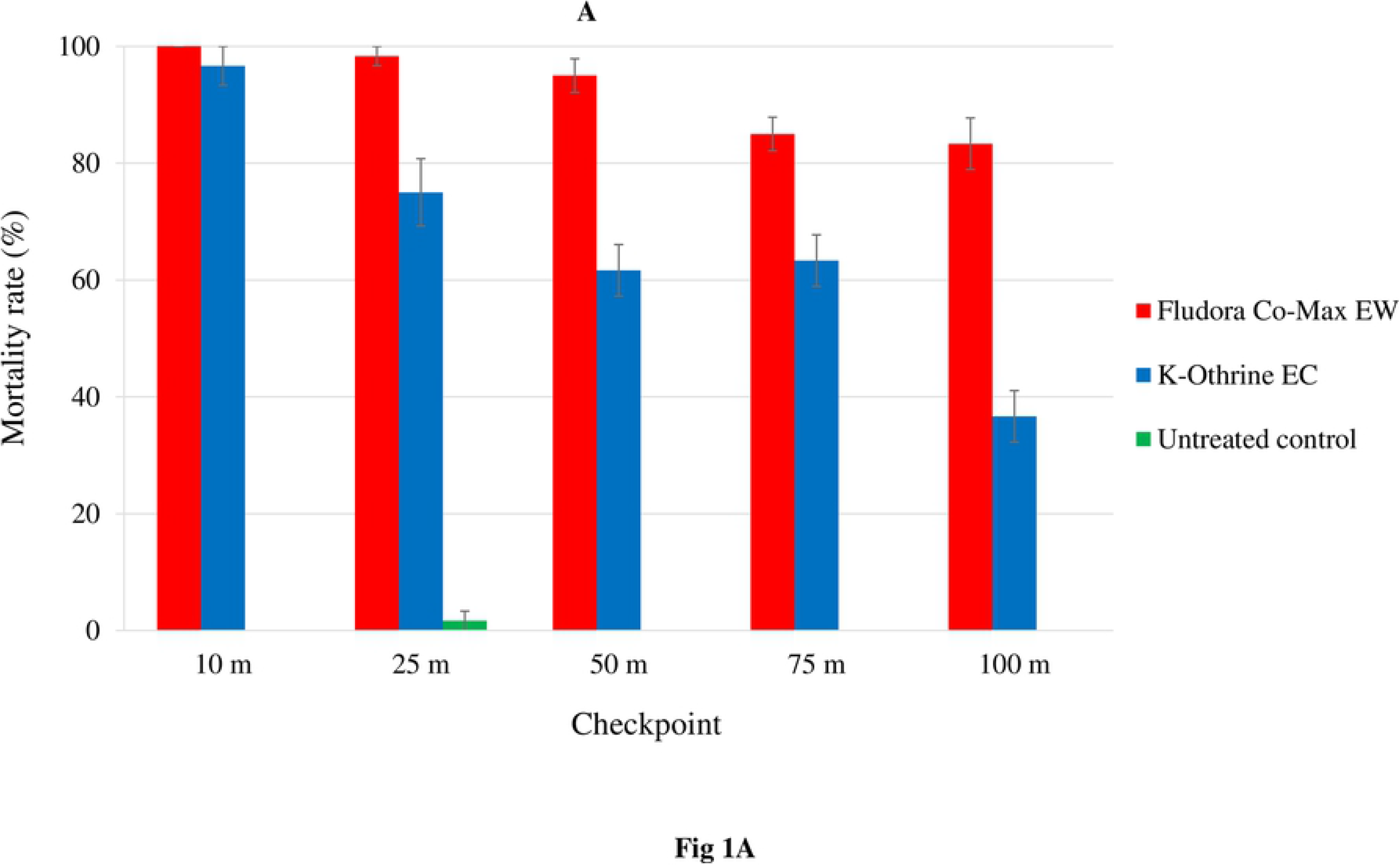
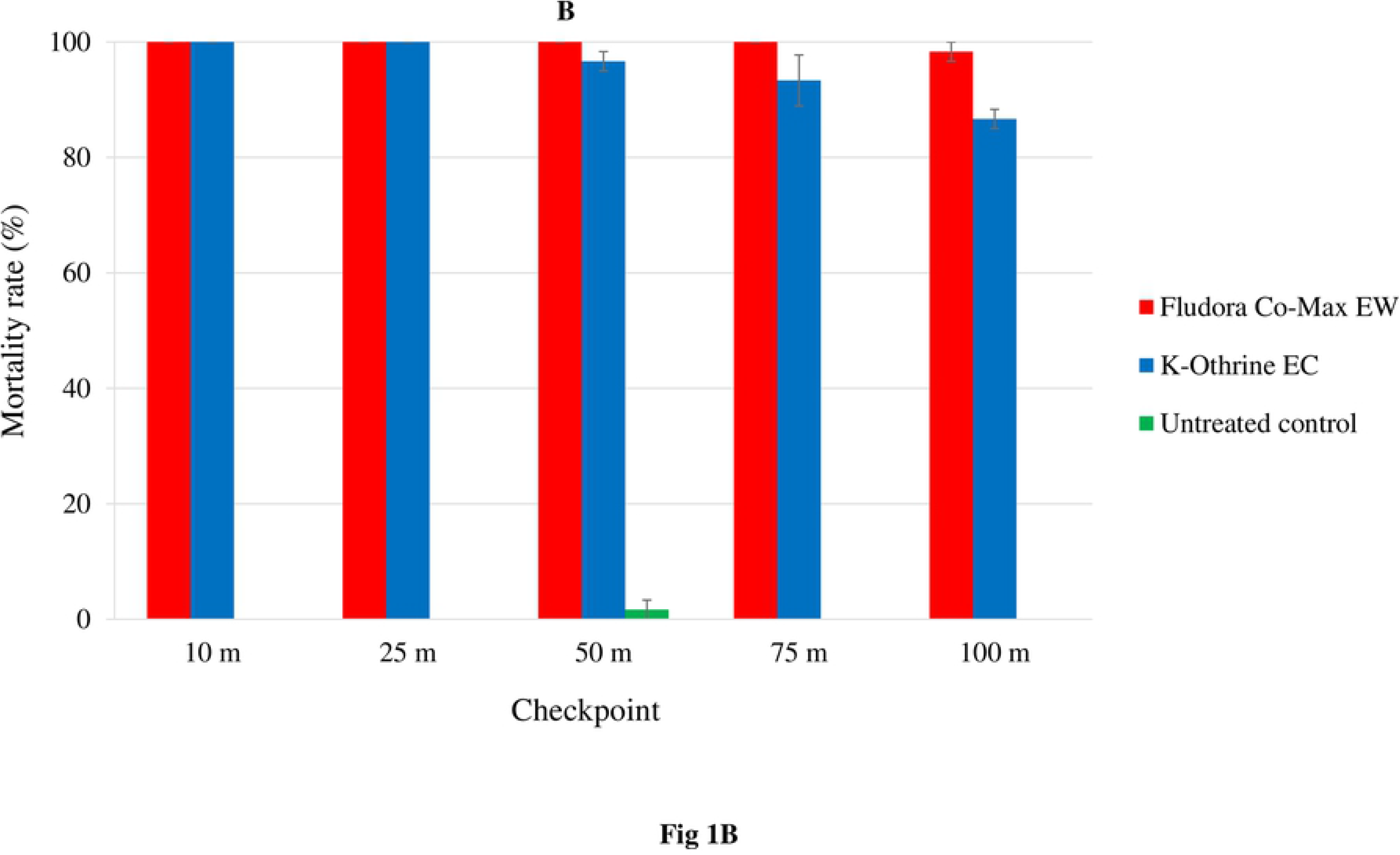
Mortality of the wild insecticide-resistant *Aedes aegypti* and *Culex quinquefasciatus* mosquito Abidjan strains exposed to outdoor ULV space sprays of Fludora Co-Max EW and K-Othrine EC. A: *Aedes aegypti*, B: *Culex quinquefasciatus*. %: percentage, m: meter, ULV: ultra-low volume. Error bars show the standard errors (SE) of the mean.

• For *Ae. aegypti*, Fludora Co-Max EW ULV space spray induced high mortality rates which varied only from 100.0 ± 0.0% at 10 m to 83.3 ± 4.4% at 100 m, and only the mortality rate at 100 m was significantly lower compared with the other distance checkpoints (p < 0.05). Conversely, K-Othrine ULV provided the high mortality of 96.7 ± 3.3% at 10 m that rapidly declined to 36.7 ± 4.4% at 100 m, and mortality rates from 50 to 100 m were was significantly lower compared with 10 and 25 m (all p < 0.05). From 25 to 100 m, Fludora Co-Max EW induced significantly higher mortality rates compared with K-Othrine EC at each distance checkpoint (p < 0.05). Fludora Co-Max EW mortality rates were always in excess of 80%, exceeding 90% between 10 m (100.0 ± 0.0%) and 25 m (91.7 ± 7.6%) while K-Othrine EC mortality was lower than 80% from 25 to 100 m.

• For *Cx. quinquefasciatus*, the mortality rates with outdoor ULV space spray ranged between 100.0 ± 0.0% at 10 m and 98.3 ± 1.7% at 100 m for Fludora Co-Max EW and between 100.0 ± 0.0% at 10 m and 86.7 ± 1.7% at 100 m for K-Othrine EC. While there was no significant reduction in Fludora Co-Max EW mortality from 10 to 100 m (χ^2^ = 4.00; df = 4; p = 0.406), K-Othrine EC mortality significantly decreased from 10 to 100 m (χ^2^ = 9.91; df = 4; p < 0.05). At 100 m, the mortality rates were significantly higher for Fludora Co-Max EW (98.3 ± 1.7%) compared with K-Othrine EC (86.7 ± 1.7%) (χ^2^ = 4.09; df = 1; p < 0.05). Overall, the mortality rates were higher than 90% for Fludora Co-Max EW at any distance checkpoint, while K-Othrine EC mortality was lower than 90% at 100 m.

### Outdoor TF efficacy

#### Knockdown

At 0 minutes post-exposure, Fludora Co-Max EW and K-Othrine EC outdoor TF space sprays provided, respectively, knockdown rates estimated at 95.0-71.0% and 71.7-36.7% in *Ae. aegypti*, and 98.3-28.3% and 66.7-16.7% in *Cx. quinquefasciatus* from 10 to 100 m (S3 Table). At 60 minutes post-exposure, the respective knockdown rates of Fludora Co-Max EW and K-Othrine EC were estimated 100.0-85.0% and 85.0-36.7% in *Ae. aegypti*, and 100.0-51.7% and 83.3-63.3% in *Cx. quinquefasciatus* (S3 Table). Fludora Co-Max EW thus caused more rapid and higher knockdown effects and up to longer distance compared with K-Othrine EC.

#### Mortality

With outdoor TF space sprays, Fludora Co-Max EW and K-Othrine EC overall mortality rates were, respectively, 73.3 ± 5.5% and 57.0 ± 5.2% in *Ae. aegypti*, and 96.3 ± 2.1% and 86.3 ± 3.1% in *Cx. quinquefasciatus* (S4 Table). Fludora Co-Max EW induced statistically higher overall mortality rates compared with K-Othrine EC in *Ae. aegypti* (F = 4.62; df = 1; p < 0.05) and in *Cx. quinquefasciatus* (χ^2^ = 7.32; df = 1; p < 0.05). Only Fludora Co-Max EW mortality rate in *Cx. quinquefasciatus* was above 90%.

The mortality rates varied significantly among the distance checkpoints (Figure 2).

**Figure 2:**
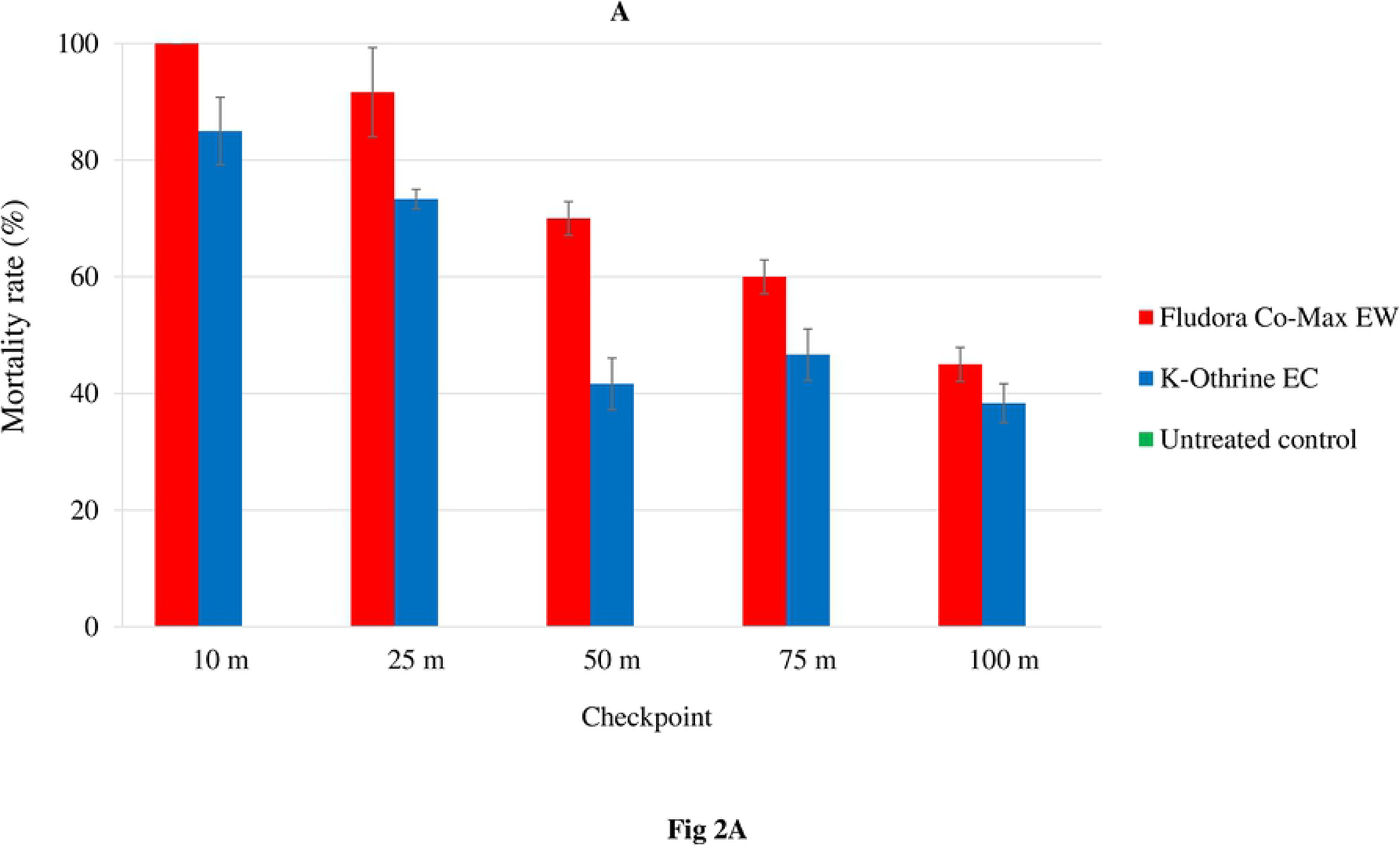
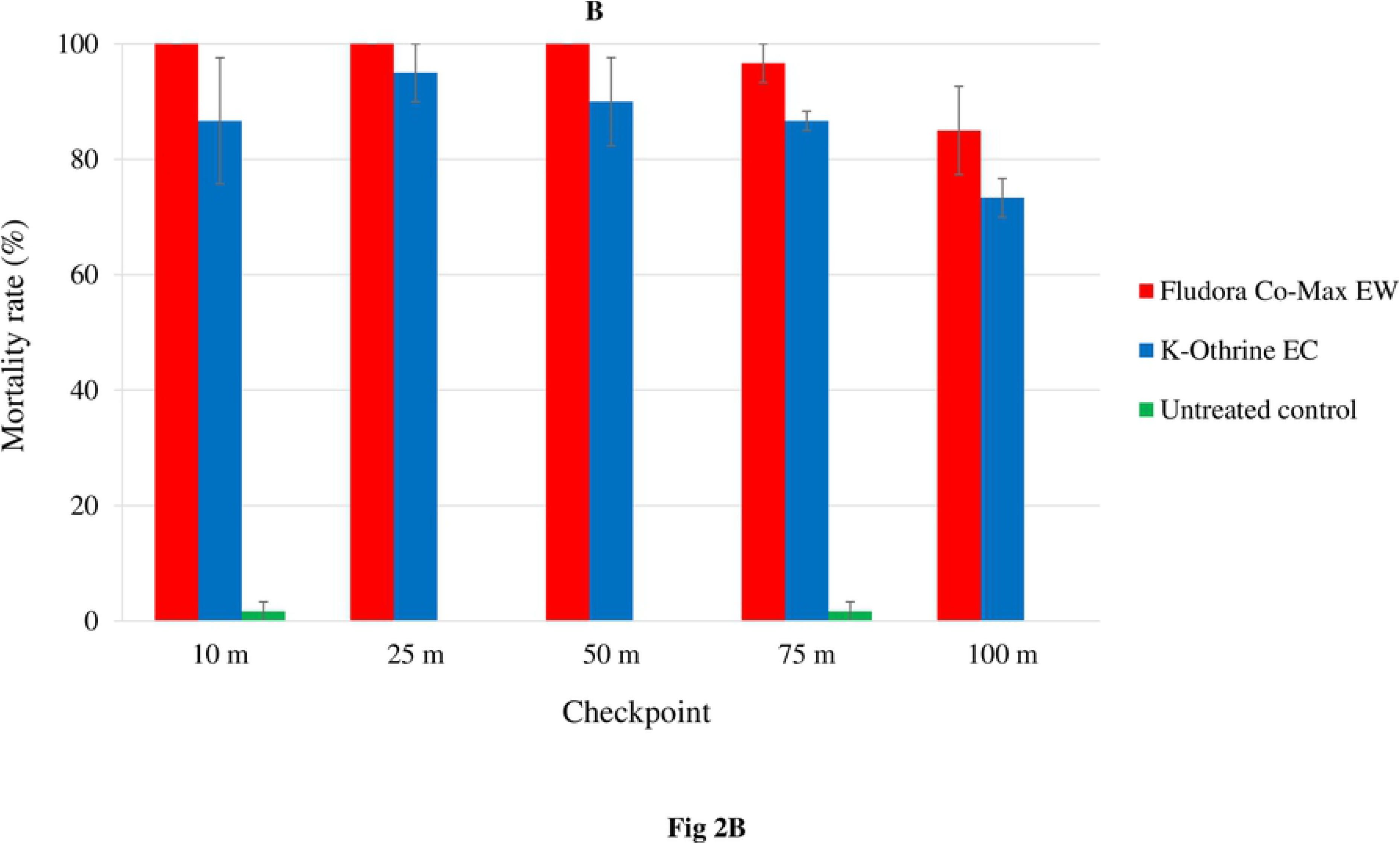
Mortality of the wild insecticide-resistant *Aedes aegypti* and *Culex quinquefasciatus* mosquito Abidjan strains exposed to outdoor TF space sprays of Fludora Co-Max EW and K-Othrine EC. A: *Aedes aegypti*, B: *Culex quinquefasciatus*. %: percentage, m: meter, TF: thermal fogging. Error bars show the standard errors (SE) of the mean.

• In *Ae. aegypti*, outdoor TF space sprays of Fludora Co-Max EW caused high mortality with rates of 100.0 ± 0.0% at 10 m and 91.7 ± 7.6% at 25 m, and the mortality significantly decreased from 70.0 ± 2.9% at 50 m to 45.0 ± 2.9% at 100 m (F = 57.34; df = 4; p < 0.001). K-Othrine mortality rates declined suddenly from 85.0 ± 5.8% at 10 m to 38.3 ± 3.3% at 100 m, with significant differences between the checkpoints (F = 25.27; df = 4; p < 0.001). Fludora Co-Max EW outdoor TF spray resulted in higher mortality compared with K-Othrine EC with significant differences at 25 and 50 m (all p < 0.05), but without statistical differences at 10, 75 and 100 m (all p > 0.05). Fludora Co-Max EW mortality rates were above 90% at 10 and 25 m, whereas K-Othrine mortality rates were below 90% at all the checkpoints.

• In *Cx. quinquefasciatus*, outdoor TF space sprays of Fludora Co-Max EW and K-Othrine EC induced high mortality. The mortality rates that decreased, without significant differences, from 100 ± 0.0% at 10 m and 85.0 ± 7.6% at 100 m for Fludora Co-Max EW (χ^2^ = 6.71; df = 4; p = 0.1520) and from 95.0 ± 5.0% at 25 m to 73.3 ± 3.3% at 100 m for K-Othrine EC (F = 1.49; df = 4; p = 0.2776). The mortality rates were substantially higher for Fludora Co-Max EW compared with K-Othrine EC, but lacked statistical difference (χ^2^ = 16.71; df = 9; p = 0.0535). Mortality rates were higher than 90% for both Fludora Co-Max EW and K-Othrine EC up to 100 m and 75 m checkpoints, respectively.

### Indoor ULV efficacy

#### Knockdown

At 0 minutes after exposure, indoor ULV space sprays of Fludora Co-Max EW and K-Othrine EC induced respective knockdown rates of 99.6-100.0% and 89.2-93.3% in *Ae. aegypti*, and 92.1-100.0% and 75.0-92.1% in *Cx. quinquefasciatus* (S5 Table). At 60 minutes post-exposure, Fludora Co-Max EW and K-Othrine EC knockdown rates were estimated at 100.0% and 97.5-99.2% for *Ae. aegypti*, and 99.2-100.0% and 97.5-100.0% for *Cx. quinquefasciatus* (S5 Table). Thus, knockdown effects were higher and faster with Fludora Co-Max EW than K-Othrine EC.

#### Mortality

Indoor UVL space sprays of Fludora Co-Max EW and K-Othrine EC resulted in the overall high mortality rates of 100.0 ± 0.0% and 97.8 ± 0.6% in *Ae. aegypti*, respectively (S6 Table). Fludora Co-Max EW induced statistically higher mortality rates compared with K-Othrine EC (χ^2^ = 4.918; df = 1; p < 0.05). Both insecticide products induced overall mortality rates of 100.0 ± 0.0% in *Cx. quinquefasciatus* (S6 Table). Therefore, the mortalities with both Fludora Co-Max EW and K-Othrine EC were largely above 90% in both mosquito species.

The mortality rates were very high at all position level checkpoints, but varied significantly among the checkpoints (Figure 3).

**Figure 3:**
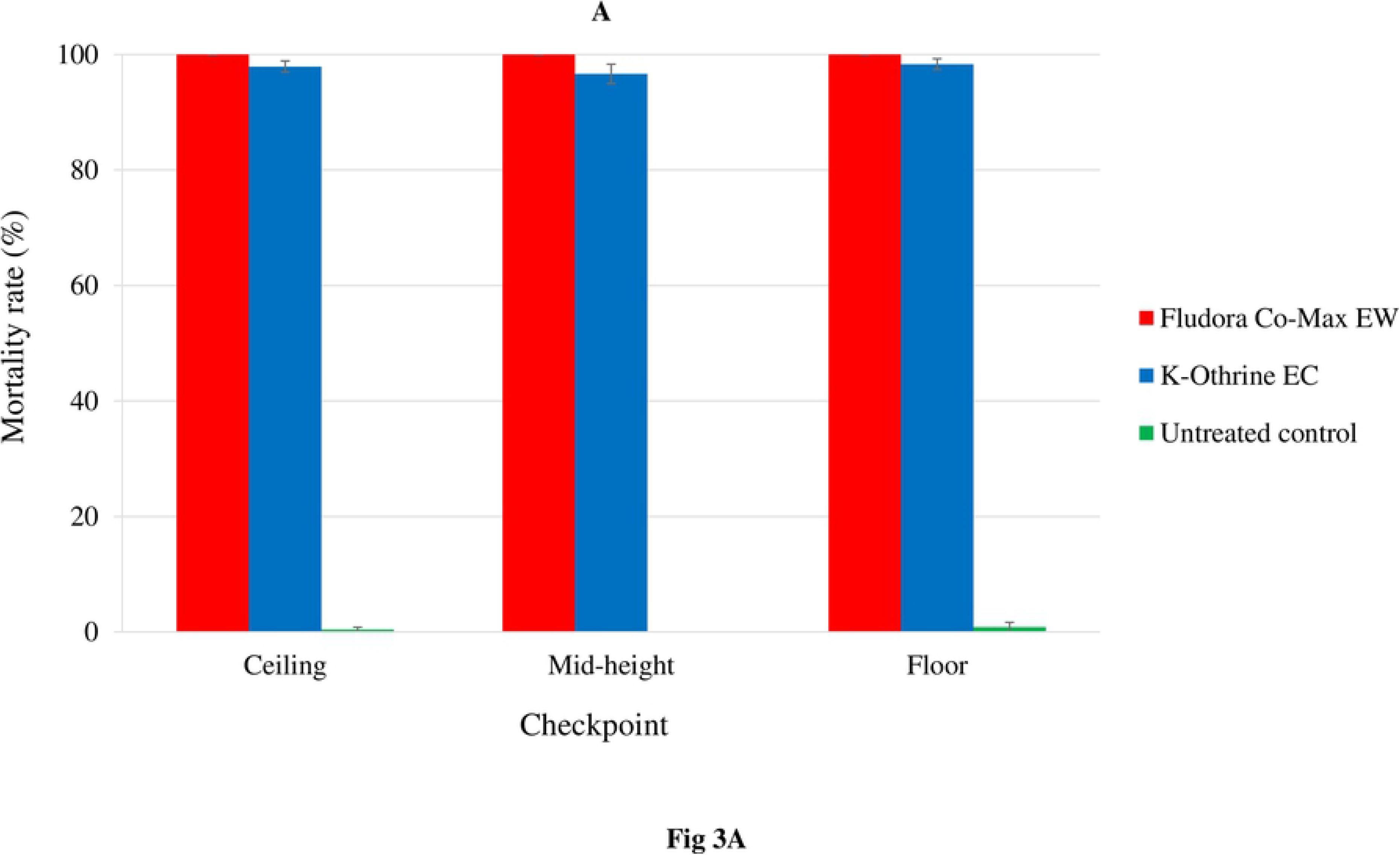
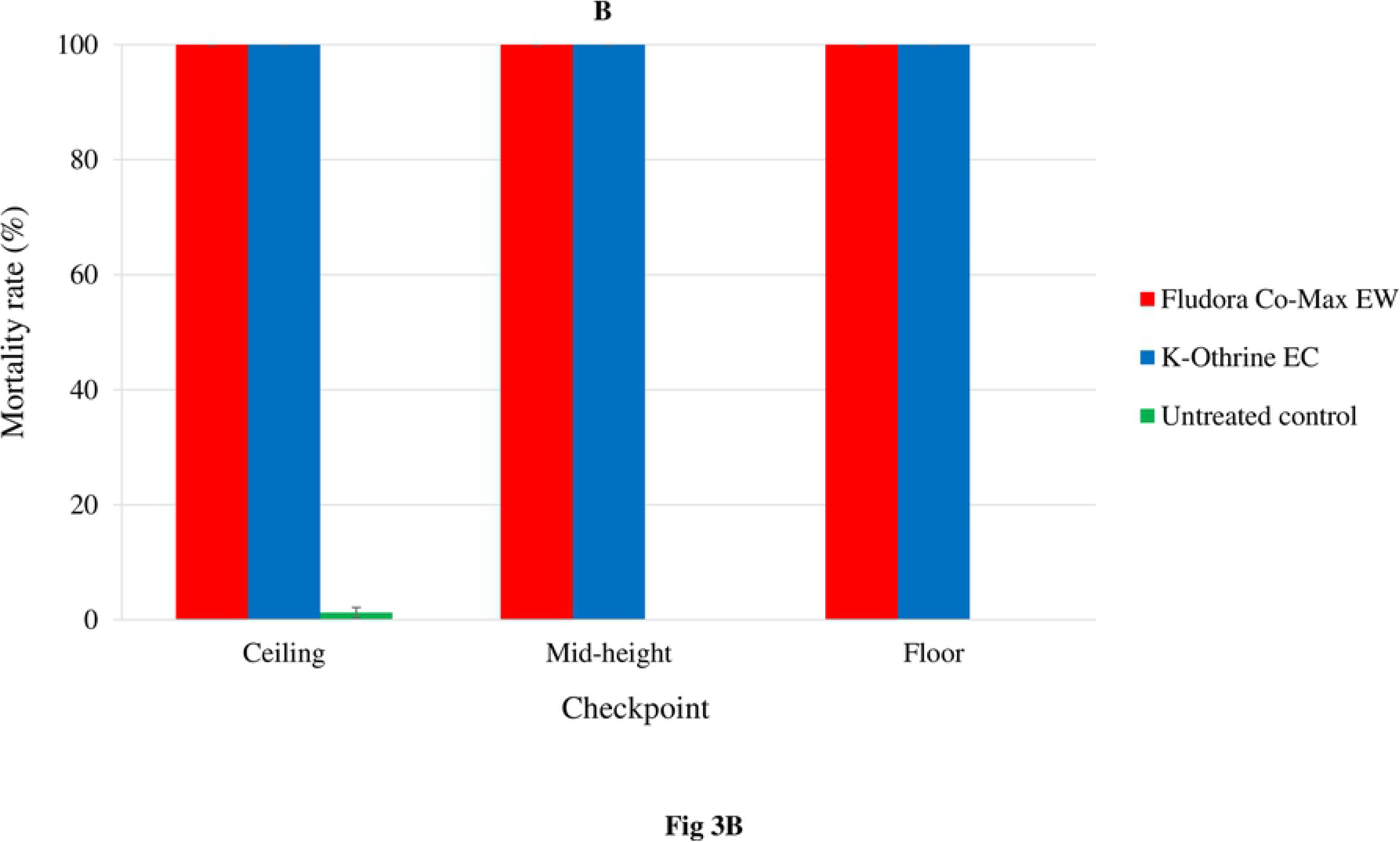
Mortality of the wild insecticide-resistant *Aedes aegypti* and *Culex quinquefasciatus* mosquito Abidjan strains exposed to indoor TF space sprays of Fludora Co-Max EW and K-Othrine EC. A: *Aedes aegypti*, B: *Culex quinquefasciatus*. %: percentage, m: meter, ULV: ultra-low volume. Error bars show the standard errors (SE) of the mean.

• In *Ae. aegypti* Fludora Co-Max EW induced 100.0 ± 0.0% mortality at any cage level position checkpoints. K-Othrine EC mortality rates were high as well, and varied between 98.3 ± 0.9% at the floor and 96.7 ± 1.7% at the mid-height. Compared with K-Othrine EC, Fludora Co-Max EW mortality rates were statistically higher at the ceiling (χ^2^ = 4.57; df = 1; p < 0.05), but not significantly different at the mid-height level (χ^2^ = 4.57; df = 1; p = 0.0578) and floor (χ^2^ = 3.27; df = 1; p = 0.0704).

• In *Cx. quinquefasciatus*, both Fludora Co-Max EW and K-Othrine EC produced very high mortality rates of 100.0 ± 0.0% at all position level checkpoints in house.

### Indoor TF efficacy

#### Knockdown

At 0 minutes after exposure, Fludora Co-Max EW and K-Othrine EC indoor TF space spraying induced high knockdown effects, with respective knockdown rates of 100.0% and 89.2-92.1% in *Ae. aegypti*, and 99.2-100.0% and 78.3-95.8% in *Cx. quinquefasciatus* (S7 Table). At 60 minutes post-exposure, Fludora Co-Max EW and K-Othrine EC knockdown rates were, respectively, of 100.0% and 98.8-100.0% for *Ae. aegypti,* and 100.0% and 93.8-100.0% for *Cx. quinquefasciatus* (S7 Table). Overall, Fludora Co-Max EW had faster and higher knockdown effects compared with K-Othrine EC.

#### Mortality

With indoor TF space sprays, overall mortality rates in *Ae. aegypti* were 100.0% ± 0.0% for Fludora Co-Max EW and 99.3 ± 0.3% for K-Othrine EC (S8 Table). Fludora Co-Max EW performed significantly better than K-Othrine EC (χ^2^ = 4.21; df = 1; p < 0.05). In *Cx. quinquefasciatus*, both Fludora Co-Max EW and K-Othrine EC provided each very high mortality rates of 100.0 ± 0.0% (S8 Table). The overall mortality rates with both insecticide products were above 90% in the two mosquito species.

The mortality rates varied significantly among the position level checkpoints (Figure 4).

**Figure 4:**
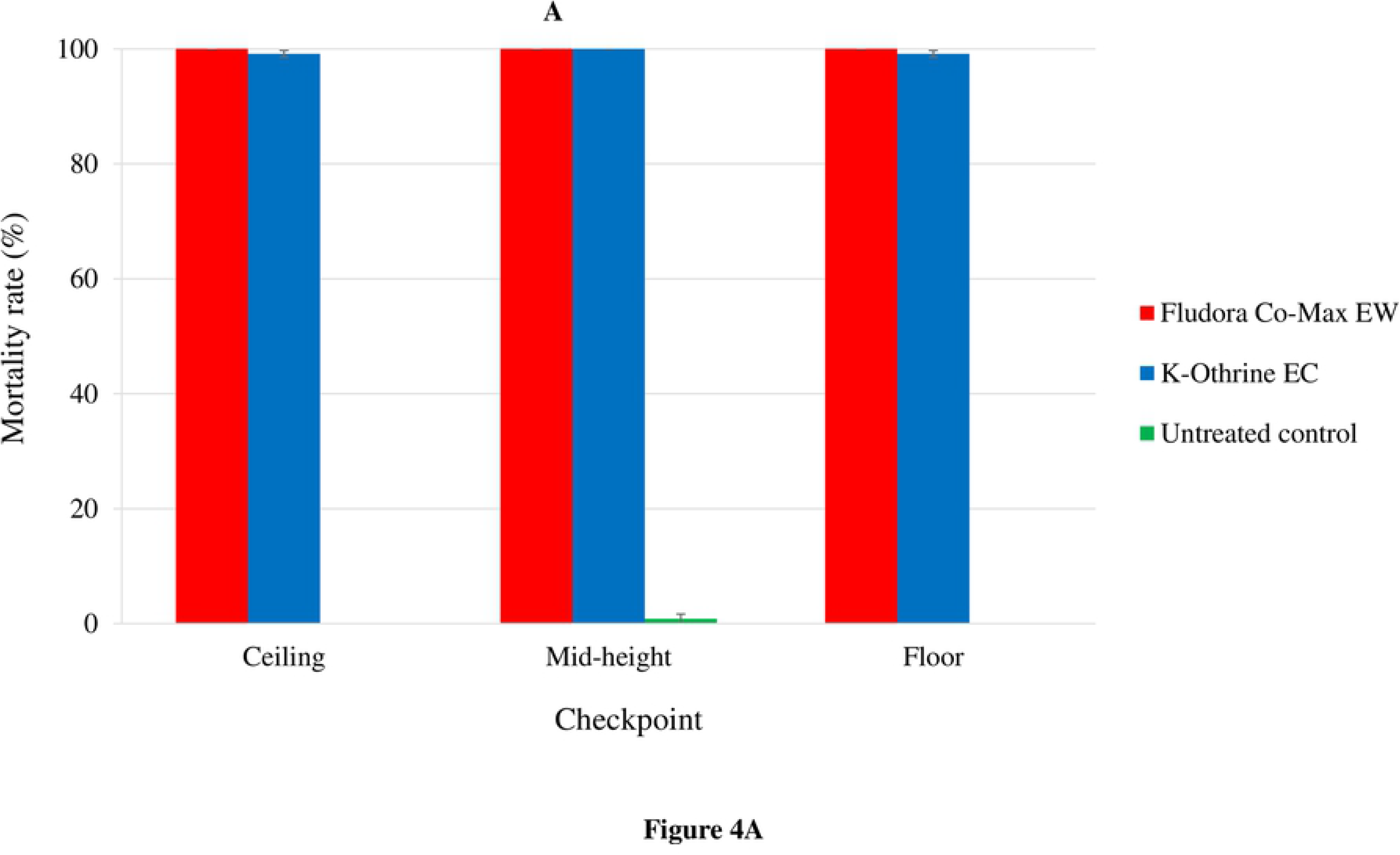
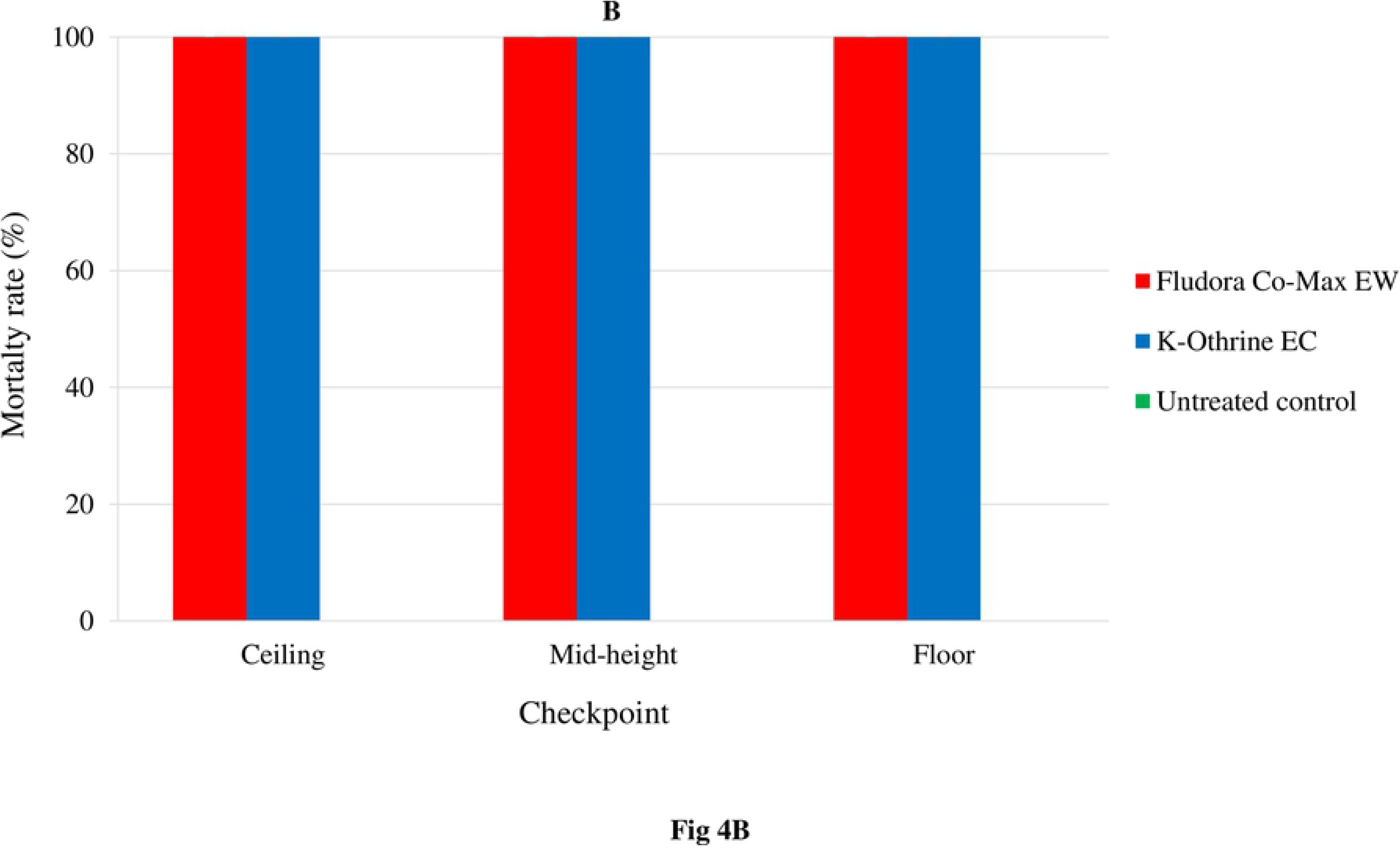
Mortality of the wild insecticide-resistant *Aedes aegypti* and *Culex quinquefasciatus* mosquito Abidjan strains exposed to indoor TF space sprays of Fludora Co-Max EW and K-Othrine EC. A: *Aedes aegypti*, B: *Culex quinquefasciatus*. %: percentage, m: meter, TF: thermal fogging. Error bars show the standard errors (SE) of the mean.

• For *Ae. aegypti* Fludora Co-Max EW always induced 100.0% mortality rate at any level checkpoints, while K-Othrine EC caused high mortality rates that varied from 99.2 ± 0.6 at the ceiling and the floor to 100.0 ± 0.0% at the mid-height level. However, there was no significant difference between and K-Othrine EC mortalities regardless the level checkpoints (χ^2^ = 1.12; df = 1; p = 0.5725). No significant differences were observed between Fludora Co-Max EW and K-Othrine EC mortality rates regardless the level checkpoints in the house (p > 0.05). However, Fludora Co-Max EW performed better than K-Othrine EC.

• For *Cx. quinquefasciatus*, both Fludora Co-Max EW and K-Othrine EC products, regardless of application method, scored mortality rates of 100% ± 0.0% at all position checkpoints.

### Fludora Co-Max EW space spray efficacy: ULV *versus* TF

For outdoor space sprays, Fludora Co-Max EW resulted in overall mortality rates of 92.3 ± 2.1% with ULV and 73.3 ± 5.5% with TF in *Ae. aegypti*, and of 99.7 ± 0.3% with ULV and 96.3 ± 2.2% with TF in *Cx. quinquefasciatus* (S2, 4, 6 and 8 Tables). Compared with TF, ULV showed higher mortality rates with significant difference for *Ae. aegypti* (χ^2^ = 5.52; df = 1; p < 0.05), but not for *Cx. quinquefasciatus* (χ^2^ = 1.34; df = 1; p = 0.2469). Especially, *Ae. aegypti* mortality rates were high and varied between 100.0 ± 0.0% and 83.3 ± 4.4% from 10 to 100 m for ULV, with a marked decline from 100.0 ± 0.0% at 10 m to 45.0 ± 2.9% at 100 m when using TF (Figure 5). *Ae. aegypti* mortality rates were statistically higher with ULV compared with TF at 50, 75 and 100 m (F = 33.32; df = 5; p < 0.001).

**Figure 5:**
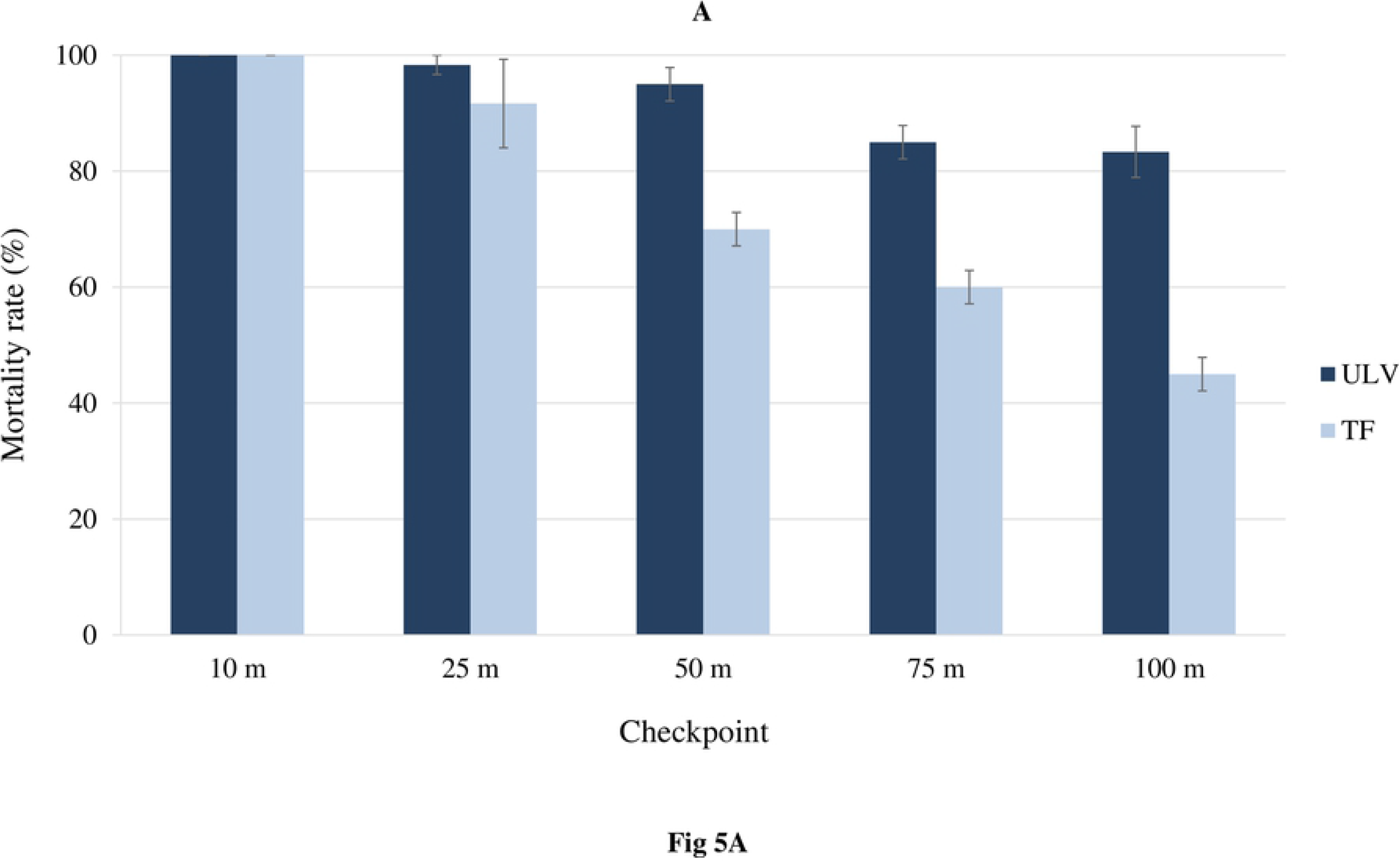
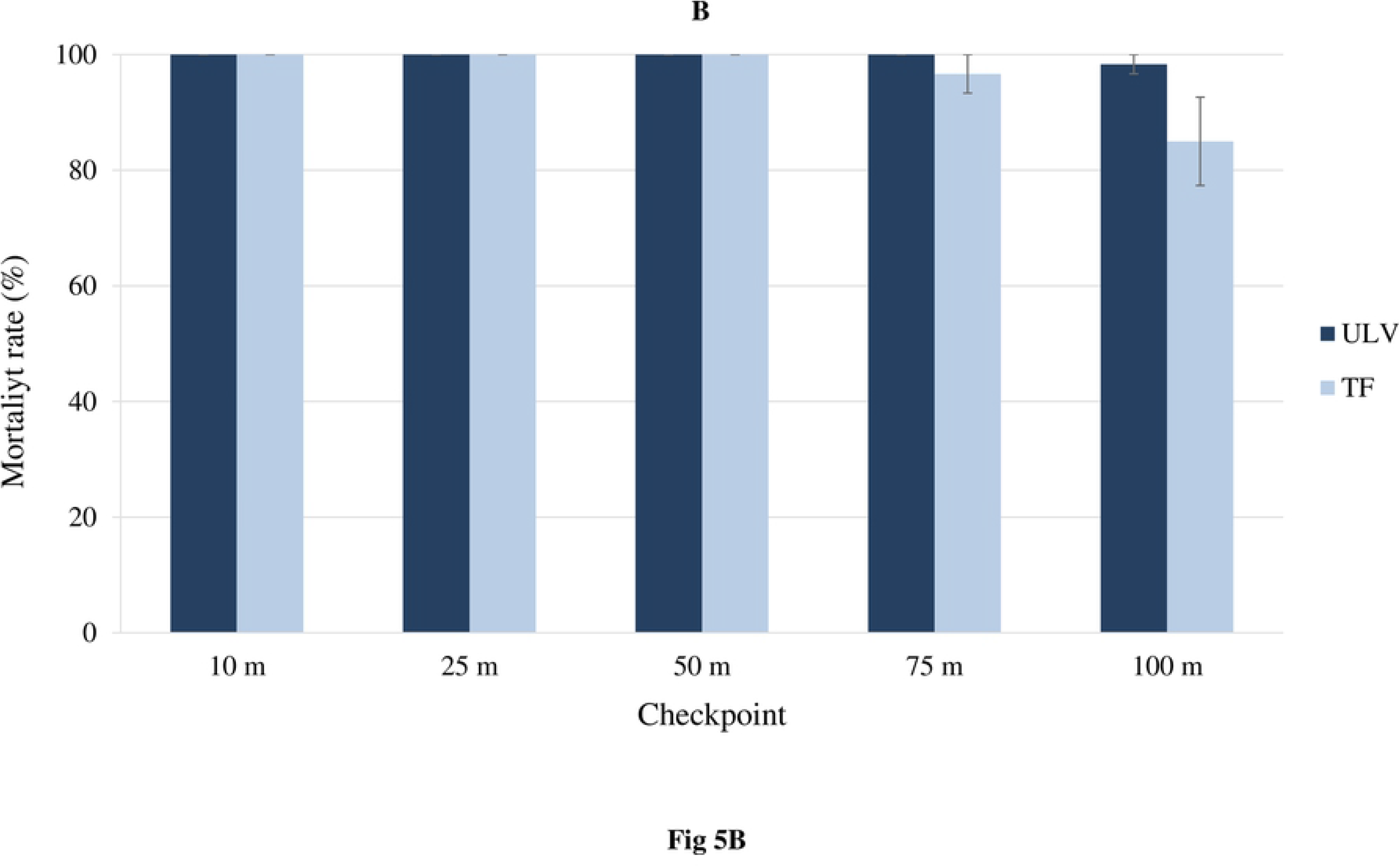
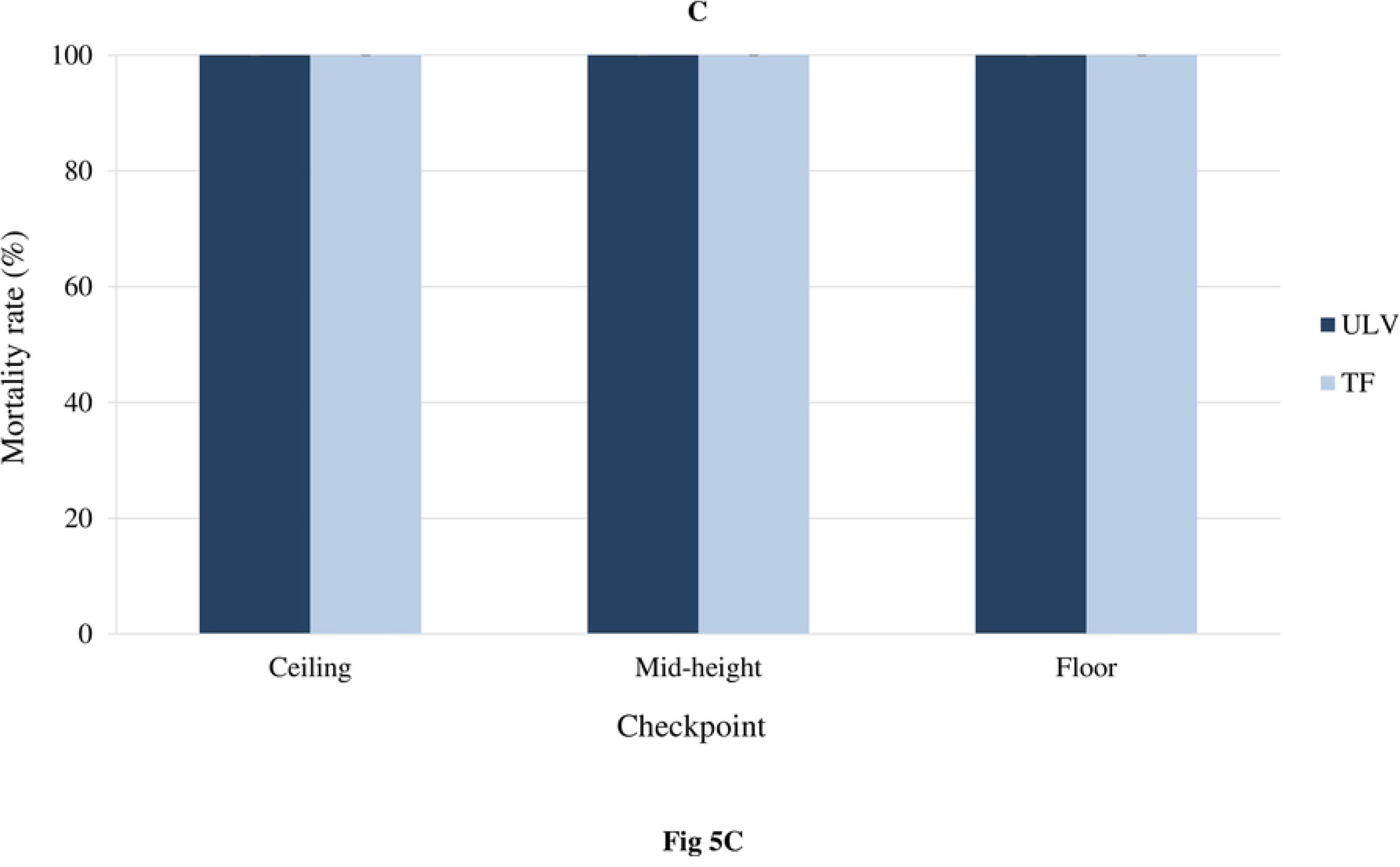
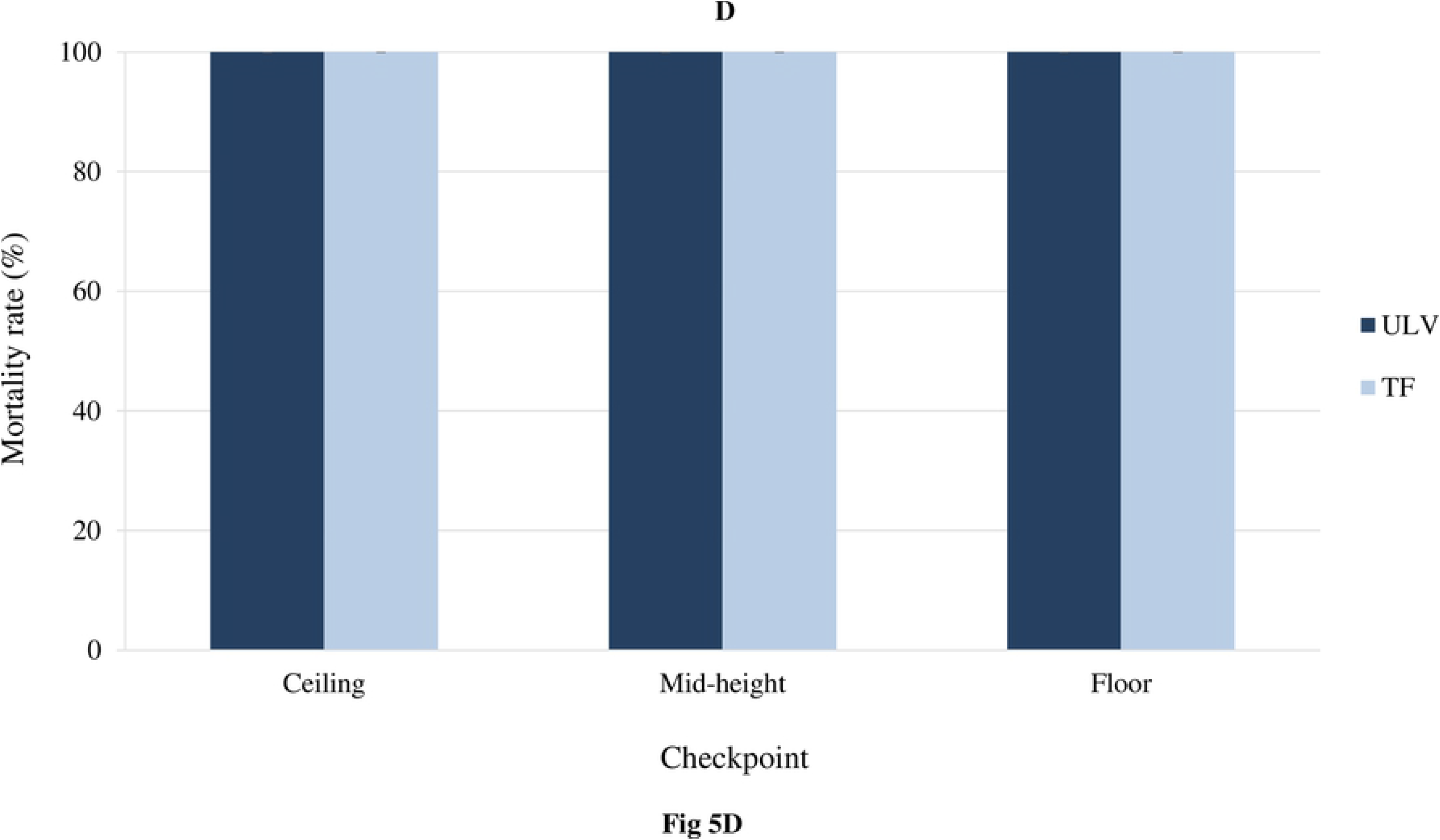
Comparison between ULV and TF space spray applications of Fludora Co-Max EW against the wild insecticide-resistant *Aedes aegypti* and *Culex quinquefasciatus* mosquito Abidjan strains. A: outdoor ULV, B: outdoor TF, C: indoor ULV, D: indoor TF. %: percentage, m: meter, TF: thermal fogging, ULV: ultra-low volume. Error bars show the standard errors (SE) of the mean.

For Fludora Co-Max EW indoor space sprays, ULV and TF both induced 100.0 ± 0.0% mortality in both *Ae. aegypti* and *Cx. quinquefasciatus* overall and at any level checkpoints in the house (Figure 5). Thus, no differences were found between ULV and TF mortality rates in either in *Ae. aegypti* and nor in *Cx. quinquefasciatus* indoors.

#### Perceived side effects

No negative side effects were reported among the sprayers, the operators and the occupants of the sprayed house during and after spraying.

## Discussion

The current phase II study aimed to evaluate and compare the outdoor and indoor space spray efficacy of Fludora Co-Max EW (candidate product) and K-Othrine EC (positive reference control) against caged adult *Ae. aegypti* and *Cx. quinquefasciatus* mosquitoes from Abidjan, Côte d’Ivoire using cold ULV and TF applications through small-scale field trials. To our knowledge, this study represents one of the first small-scale trials evaluating the space spray efficacy of an insecticide against *Aedes* and *Culex* mosquitoes in sub-Saharan Africa. The results revealed that Fludora Co-Max EW was effective against wild insecticide-resistant *Ae. aegypti* and *Cx. quinquefasciatus*, and performed better than K-Othrine EC in the field.

Overall, the current semi-field evaluations showed that ULV and TF space spray of Fludora Co-Max EW outdoors and indoors effected high knockdown and mortality rates (>90%) in both *Ae. aegypti* and *Cx. quinquefasciatus* Abidjan strains, and the values were higher or equal compared with K-Othrine EC. Additionally, knocking down effects in both strains were faster with Fludora Co-Max EW than with K-Othrine EC. The reduction in *Aedes* and *Culex* Abidjan strain susceptibility and mortality to K-Othrine EC (deltamethrin being the only active ingredient) may be due to their resistance to pyrethroids [16–18], as reported on the island of Martinique in Marcombe et al. [27]. However, the high effectiveness of Fludora Co-Max EW against the same insecticide-resistant mosquito strains may be explained by the mixture of two active ingredients, namely transfluthrin and flupyradifurone. Indeed, the Fludora Co-Max EW formulation has two active ingredients, the butenolide flupyradifurone and the pyrethroid transfluthrin, representing two unrelated modes of action chosen for their complementary action, making it harder for resistance development, and helps to control pyrethroid-resistant mosquitoes [28, 29]. Flupyradifurone belongs to the group of nicotinic acetylcholine receptor (nAChR) competitive modulators that bind to the acetylcholine site on nAChRs, causing a range of symptoms from hyper-excitation to lethargy and paralysis [28, 29]. Transfluthrin belongs to the group of sodium channel modulators that keep sodium channels open, causing hyper-excitation as well and, in some cases, nerve block [29]. Therefore, as suggested by WHO [26], an overall adult vector mortality of at least 90% is considered effective, thus recommending for Fludora Co-Max EW for the control of *Ae. aegypti* and *Cx. quinquefasciatus* outdoors and indoors, if even in areas with high insecticide resistance. Generally, both mosquito species mortality was higher in all indoor and outdoor cages from the ULV spraying when compared with the TF spraying, but this difference was only significant outdoors. Moreover, specific differences were found in Fludora Co-Max EW and K-Othrine EC efficacy according to the methods, and distance or position checkpoints.

The outcomes of this semi-field study demonstrated that outdoor ULV space spraying of Fludora Co-Max EW produced in *Ae. aegypti* mortality rates maximal (100%) at up to 25 m, decreasing, while remaining above 90% up to 50 m, and 80% up to 100 m. In contrast, the mortality rates of K-Othrine EC rapidly dropped down below 80% from 25 m and continued to decrease at up to 100 m. In *Cx. quinquefasciatus,* Fludora Co-Max EW mortality rates varied between 100% at 10 m and 98% at 100 m, while K-Othrine EC induced mortality rates were 100% at 10 m and decreased to 87% at 100 m. The capacity of Fludora Co-Max EW to kill mosquitoes over long distance is consistent with the results reported by Farooq et al. [10] where ULV application of Kontrol (permethrin and piperonyl butoxide) produced 100% mortality in *Ae. aegypti* outdoors in Starke, Florida, USA. However, Kontrol contains a synergist and the distance checkpoints were shorter (25-30 m) [10] compared with the Fludora Co-Max EW effect distance checkpoints (10-100 m). Moreover, Fludora Co-Max EW kill mosquitoes up to significantly longer distance compared to K-Othrine EC. Indeed, Fludora Co-Max EW incorporates FFAST anti-evaporant technology, and during formation of the spray droplets, the anti-evaporant agent forms a protective skin around the droplets that slows down evaporation events. This process ensures the optimal droplet size is maintained longer, prolonging maximal efficacy duration, allowing Fludora Co-Max EW to achieve higher mortality up to 100 m.

This study revealed that outdoor TF space sprays of Fludora Co-Max EW and K-Othrine EC displayed high knockdown and mortality effects in *Ae. aegypti* and *Cx. quinquefasciatus* showing similar tendencies observed for ULV, but with more pronounced decline for TF and K-Othrine EC. For instance, Fludora Co-Max EW mortality rates were above 90% only in *Cx. quinquefasciatus*. Additionally, Fludora Co-Max EW mortality rates were 90-100% from 10 to 25 m, and significantly decreased to 70-45% from 50 to 100 m in *Ae. aegypti* and decreased from 100% at 10 m to 85% at 100 m in *Cx. quinquefasciatus*. Such lower mortality with outdoor TF (85%) compared to outdoor ULV (100%) was observed with Kontrol in Stake, USA [10]. The reduction in Fludora Co-Max EW mortality with TF outdoors might be attributed to thermal effect (i.e. high temperature) that may have reduced the size of spray droplets. This may have allowed the wind to change the direction of droplets and prevent insecticide to reach and kill caged mosquitoes from a longer distance (Fludora Co-Max EW indoor TF induced 100% mortality in the same *Ae. aegypti* and *Cx. quinquefasciatus* strains inside of a closed house). Indeed, sudden changes of wind movement and droplet size can affect the homogeneity of droplet depositions, and thus play an important role in drifting [30].

The results of this study exhibited that indoor ULV applications of Fludora Co-Max EW provided 100% knockdown and 100% mortality in local wild insecticide-resistant populations of *Ae. aegypti* and *Cx. quinquefasciatus*. While K-Othrine EC was able to induce high knockdown (92-100%) and mortality (98-100%), but the rates were lower than Fludora Co-Max EW. Moreover, Fludora Co-Max EW induced high mortality (∼100%) in both mosquito species at all position level checkpoints in a house. Indoor ULV applications of Fludora Co-Max EW carried out in this study were more effective than applications performed against *Ae. aegypti* using other products in other areas in America [9, 32] and Asia [33]. Harwood et al. [32] reported that indoor applications of the adulticide ULD BP-300 (synergized pyrethrins) with a backpack ULV sprayer resulted in 88-100% mortality in caged *Ae. aegypti*. Another study on indoor application of ULD BP-300 and 9M Plus2 (deltamethrin) indicated that mortality of caged mosquitoes in the middle of a room ranged from 59 to 100% for TF and 94 to 100% for ULV in Thailand, respectively [33]. However, 100% mortality was found with indoor ULV spraying of Kontrol in USA [9]. The high performance of indoor UVL spray of Fludora Co-Max EW be due to limited variations in the wind movement inside of house [9].

The outcomes of this trial highlighted that indoor TF space spray of Fludora Co-Max EW K and Othrine EC effected both 100% knockdown and 100% mortality in both *Ae. aegypti* and *Cx. quinquefasciatus* Abidjan strains. Conversely, indoor TF spray of Kontrol induced only 34% in *Ae. aegypti* in USA [9]. Here, the low impact of natural unpredictable changes in wind movement may have greatly improved Fludora Co-Max EW TF efficacy indoors [9].

The present study showed that the performance of space spray of Fludora Co-Max EW was influenced by the type of application methods used. ULV method induced higher mortality in *Ae. aegypti* and *Cx. quinquefasciatus* compared with TF method outdoors, and similar mortality (up to 100%) in both mosquito species indoors. Fludora Co-Max EW droplets from the ULV application and hence the active ingredient deposition was probably larger than from TF application, thus resulting in higher mortality effects outdoors for ULV, as observed in Kontrol by Farooq et al. [9]. Farooq et al. [9] also found that the active ingredient deposition from the ULV sprayer was significantly higher than from the thermal fogger both indoors and outdoors [9]. The high temperature (heat of 1,000 °C) with TF application that can increase the evaporation of the fogging insecticide that coupled with changes of wind speed and direction outside in an open area can negatively affect the homogeneity of droplets deposition may have probably reduced the efficacy of Fludora Co-Max EW outdoors [30, 31]. The low mortality in caged mosquitoes using TF may also be a result of overheating of the thermal fogger, which would breakdown the active ingredients much faster. The higher performances (100% mortality) of ULV and especially of TF indoors might be explained by limited effects of in the ambient conditions inside a closed house likely subjected to restricted environmental variations. Thus, ULV should be recommended for outdoor spray, and preferentially suggested for indoor spray due to logistical reason and to minimize evaporation. However, evaluations of droplet size and active ingredient deposition with ULV and TF applications outdoors and indoors are required to better understand the variations in Fludora Co-Max EW efficacy.

Finally, over being effective against natural insecticide-resistant populations of *Aedes* and *Culex* outdoors and indoors, Fludora Co-Max EW presents other key environmental and economic advantages. Indeed, the very low rates of application (1:10 for ULV and 1:100 for TF) and biodegradable formulation diluted with water reduce the risk of environmental accumulation and for non-target insects. Studies have shown that the insecticides mixed with water cause statistically higher or equal mosquito mortality compared with mineral oil, diesel oil or gasoline [32, 33]. Fludora Co-Max EW efficacy to kill large number of wild insecticide-resistant mosquitoes over longer distance (up to 100 m) can reduce the quantity and the cost of the insecticide and financial efforts, and thus save resources and work time. Additionally, no adverse events and complains were reported among the sprayers and the occupants of the sprayed house during and after the sprays, and this might increase community acceptability and adherence for a large use of Fludora Co-Max EW. This may be beneficial to the optimal coverage of application required to interrupt the transmission of arboviral diseases. Therefore, Fludora Co-Max EW appears to be a promising tool targeting adult vectors outdoors and indoors for the prevention, the control and the outbreak responses against arboviruses, even in areas with insecticide resistance. However, further validations are needed in a large-scale trial to assess the acceptability of Fludora Co-Max EW and for the operational vector control of free-flying natural populations of *Aedes* and *Culex* mosquitoes [34–37].

## Conclusions

The current small-scale field study demonstrated that outdoor and indoor space spraying of Fludora Co-Max EW effectively induces high knockdown (up to 100%) and mortality rates (up to 100%) in caged wild insecticide-resistant *Ae. aegypti* and *Cx. quinquefasciatus* strains from Abidjan, Côte d’Ivoire. While both ULV and TF space spray methods displayed similar efficacy indoors, ULV application was found more suitable in outdoor space spray situations. Overall, the knockdown and mortality effects in both mosquito species were higher with Fludora Co-Max EW compared with K-Othrine EC. Fludora Co-Max EW thus appears to be an effective tool for outdoor and indoor space sprays targeting in areas with insecticide resistance of *Ae. aegypti* and *Cx. quinquefasciatus* mosquitoes for arboviral prevention, control and outbreak responses.

## Supporting Information

**S1 Fig. Different methods used for the semi-field evaluation of the efficacy of Fludora Co-Max EW and K-Othrine against *Aedes aegypti* and *Culex quinquefasciatus* Abidjan strain in Agboville, Côte d’Ivoire.** A: outdoor ULV, B: outdoor TF, C: indoor ULV, D: indoor TF. ULV: ultra-low volume, TF: thermal fogging.

(TIF).

**S2 Fig. Dilution of Fludora Co-Max EW and K-Othrine in the filed in Agboville, Côte d’Ivoire.**

(TIF).

**S3 Fig. Outdoor trial semi-field station with 1.5 m tall poles placed at five different distance checkpoints.** 1: 10 m, 2: 25 m, 3: 50 m, 4: 75 m, 5: 100 m.

(TIF).

**S4 Fig. Cylindrical cages constructed of fine mesh fabric (nylon) with wire frame support (diameter 10 cm x height 15 cm x tapping cover 10 cm) and containing adult mosquitoes for outdoor trial.** A: front view, B: profile view.

(TIF).

**S4 Fig. Indoor trial semi-field station with mosquito cages installed at different level checkpoints in a house.** A: ceiling, B: mid-height, C: floor.

(TIF).

**S1 Table. Knockdown rate at time intervals post-application in the wild insecticide-resistant *Aedes aegypti* and *Culex quinquefasciatus* Abidjan strains exposed to outdoor ULV space spray of Fludora Co-Max EW and K-Othrine EC.**

(DOCX).

**S2 Table. Mortality of the wild insecticide-resistant *Aedes aegypti* and *Culex quinquefasciatus* Abidjan strains exposed to Fludora Co-Max EW and K-Othrine EC using outdoor ULV space spray.**

(DOCX).

**S3 Table. Knockdown rate at time intervals post-application in wild insecticide-resistant *Aedes aegypti* and *Culex quinquefasciatus* Abidjan strains exposed to outdoor TF space spray of Fludora Co-Max EW and K-Othrine EC.**

(DOCX).

**S4 Table. Mortality of the wild insecticide-resistant *Aedes aegypti* and *Culex quinquefasciatus* Abidjan strains exposed to Fludora Co-Max EW and K-Othrine EC using outdoor TF space spray.**

(DOCX).

**S5 Table. Knockdown rate at time intervals post-application in wild insecticide-resistant *Aedes aegypti* and *Culex quinquefasciatus* Abidjan strains exposed to indoor ULV space spray of Fludora Co-Max EW and K-Othrine EC.**

(DOCX).

**S6 Table. Mortality of the wild insecticide-resistant *Aedes aegypti* and *Culex quinquefasciatus* Abidjan strains exposed to Fludora Co-Max EW and K-Othrine EC using outdoor ULV space spray.**

(DOCX).

**S7 Table. Knockdown rate at time intervals post-application in wild insecticide-resistant *Aedes aegypti* and *Culex quinquefasciatus* Abidjan strains exposed to indoor TF space spray of Fludora Co-Max EW and K-Othrine EC.**

(DOCX).

**S8 Table. Mortality of the wild insecticide-resistant *Aedes aegypti* and *Culex quinquefasciatus* Abidjan strains exposed to Fludora Co-Max EW and K-Othrine EC using outdoor TF space spray.**

(DOCX).

## Acknowledgments

The authors are grateful to the administrative staff, the health authorities, the local authorities, and residents in the study areas in Abidjan and Agboville, Côte d’Ivoire and the field mosquito collection and insecticide application teams from the Centre Suisse de Recherches Scientifiques en Côte d’Ivoire and the Institut National d’Hygiène Publique in Abidjan, Côte d’Ivoire. The authors also thank Oliver Wood for his support with revising this manuscript.

## Author Contributions

**Conceptualization:** Julien Z. B. Zahouli, Sebastian Horstmann, Benjamin G. Koudou.

**Data curation:** Julien Z. B. Zahouli, Benjamin G. Koudou.

**Formal analysis:** Julien Z. B. Zahouli.

**Funding acquisition:** Julien Z. B. Zahouli, Benjamin G. Koudou.

**Investigation:** Julien Z. B. Zahouli, Jean-Denis Dibo, Laurence Yao, Fofana Diakaridia, Benjamin G. Koudou.

**Methodology:** Julien Z. B. Zahouli, Jean-Denis Dibo, Laurence Yao, Fofana Diakaridia, Sarah D. Souza, Sebastian Horstmann, Benjamin G. Koudou.

**Project administration:** Julien Z. B. Zahouli, Laurence Yao, Benjamin G. Koudou.

**Resources:** Fofana Diakaridia, Laurence Yao, Sarah D. Souza, Sebastian Horstmann, Benjamin G. Koudou,

**Supervision:** Julien Z. B. Zahouli, Sarah D. Souza, Benjamin G. Koudou.

**Visualization:** Julien Z. B. Zahouli, Benjamin G. Koudou, Sebastian Horstmann

**Writing – original draft:** Julien Z. B. Zahouli

**Writing – review & editing:** Julien Z. B. Zahouli, Sebastian Horstmann, Benjamin G. Koudou.

